# The microbes found in the honey of New York City beehives

**DOI:** 10.1101/2020.04.16.042762

**Authors:** Tallisker Weiss, Allison Mayle, Bruce Nash

## Abstract

Bees are incredibly important to the Earth’s ecosystem and provide humans with a variety of fruits and vegetables; however, due to Colony Collapse Disorder, hives are dying at an alarming rate. Colony Collapse Disorder is caused by a number of factors such as pesticides and bacteria that kill the bees. With the increase of urban beekeeping created in an effort to replenish the bee population, little is known about the microbes the bees are interacting with in New York City. This research looked at what microbes the bees came in contact with to use as a resource in identifying the differences between the neighborhoods. Two methods were used to extract the DNA, one looking at gram-positive and gram-negative bacteria and the other looking at only gram-positive. The samples were taken from around New York City, Westchester County and Pennsylvania. The Pennsylvania sample was collected from a container of honey bought from Trader Joe’s. The reason both urban and rural samples were used was to see if there is an obvious difference in the microbes found between them.

## Introduction

Urban beekeeping is a growing phenomenon, and by identifying the microbes that go into the honey, it may be possible to understand what the bees are interacting with. Bees travel approximately one-half to three miles around their hives in search for flowers to pollinate [1-3]. Therefore, one can look at the specific areas of New York City through the DNA of the microbes that are later incorporated into the honey [4].

This experiment was based on a project called Holobiont Urbanism that looked at the debris that the bees collected while foraging in New York City and other parts of the world [5]. Our experiment differed from previous ones in that it used honey to look specifically at the differences within New York City to understand the neighborhoods on a microbial scale compared with Westchester County and store-bought honey.

Samples were collected from beehives in New York City and Westchester County. This was done to compare the city samples with those found on a farm where the bees have lots of land to forage. This way, one can see the effects that the urbanization of beekeeping has on the honey. The last samples were collected from a container of honey bought from Trader Joe’s that was produced in Pennsylvania, in order to examine the microbes found in mass-produced honey and see how they differed from those found in locally sourced honey.

## Methods

The honey samples were collected directly from their hives in February 2019, and the DNA extraction and analysis took place last March. Two methods of extraction were used: the DNeasy Blood and Tissue Kit (Qiagen) and the DNeasy PowerSoil Kit (Qiagen). The DNeasy Blood and Tissue Kit only looked at the gram-positive bacteria, while the more expensive DNeasy PowerSoil Kit looked at both gram-positive and gram-negative. Honey is an incredibly viscous substance, and it was unclear how that would affect the results of each kit. It was theorized that The DNeasy Blood and Tissue Kit would not produce the same quality of results as the DNeasy PowerSoil Kit.

The protocol for the DNeasy Blood and Tissue Kit was followed using 100ul of honey in the initial microcentrifuge tube.

The protocol for the DNeasy PowerSoil Kit was followed, beginning with 1 milliliter of honey being placed in the PowerBead tube provided in the kit.

DNA from both extraction methods underwent Polymerase Chain Reaction using primers 515F and 806R to amplify the V4 region of the 16S SSU rRNA [6]. Primer mixes were prepared with cresol red loading dye, to a final concentration of 0.26 picomoles/μL of each primer. Then, 2 microliters of DNA were added to 23 microliters of the primer mix and an illustra PuReTaq Ready-To-Go PCR bead (GE Lifesciences) before the samples were placed into a thermocycler. Thermocycling was as described by the Earth Microbiome Project [7]. The samples were then sent to Genewiz for Amplicon-EZ Next Generation Sequencing on an Illumina platform using paired-end 250bp reads.

DNA sequence analysis was performed using the DNA Subway Purple line for microbiome analysis with the Qiime2 toolkit (DNASubway.cyverse.org).

## Results

Cosmos ID was used to create krona graphs for Figures 1-12 out of the DNA sequences that were found. The data was also run through the Purple line on DNA Subway to create Figure 13 and Tables 1 and 2.

**Figure 1:**
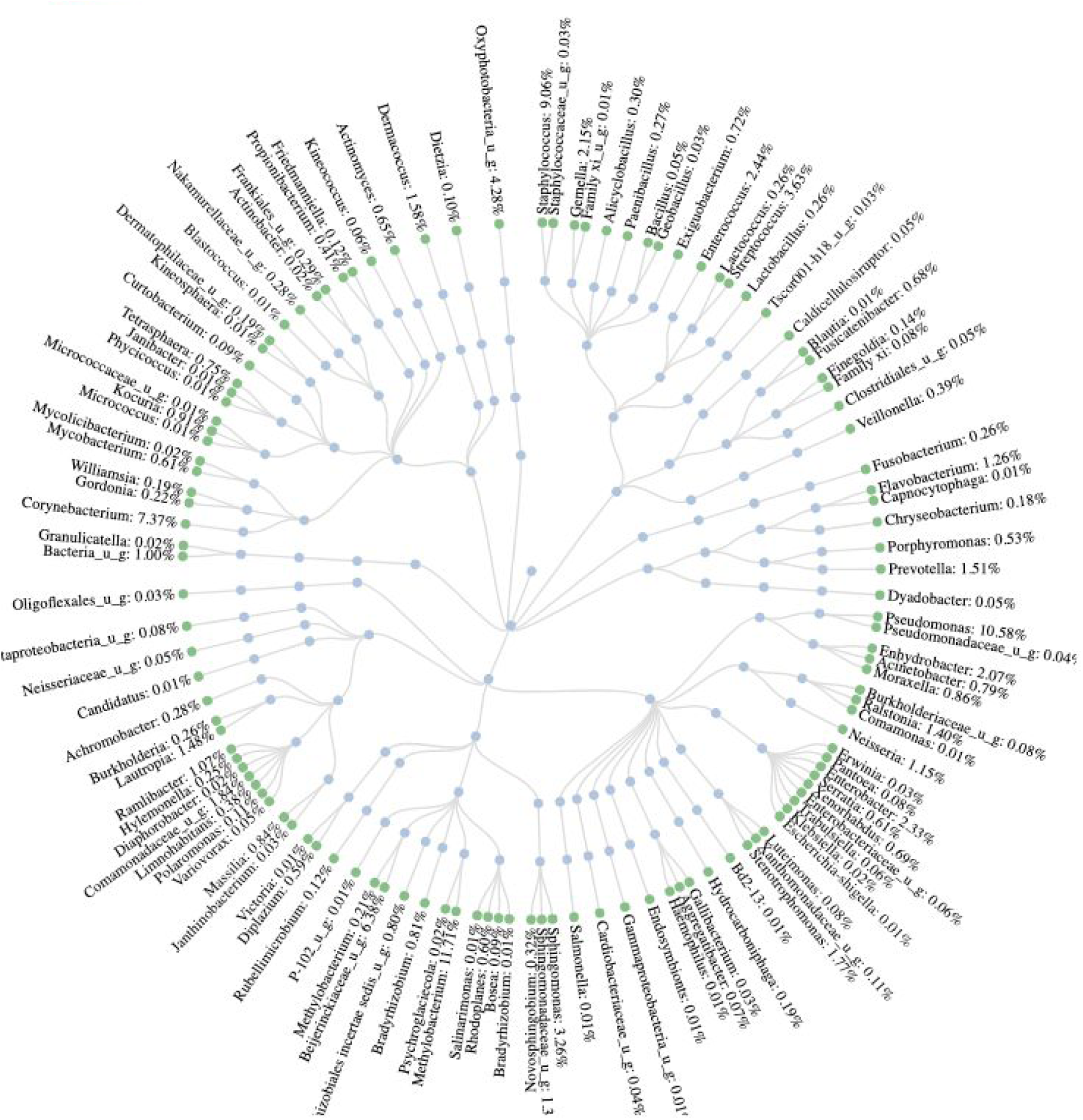
Krona graph of the microbes found in the 86 4th Ave. sample using the Soil Kit.

**Figure 2:**
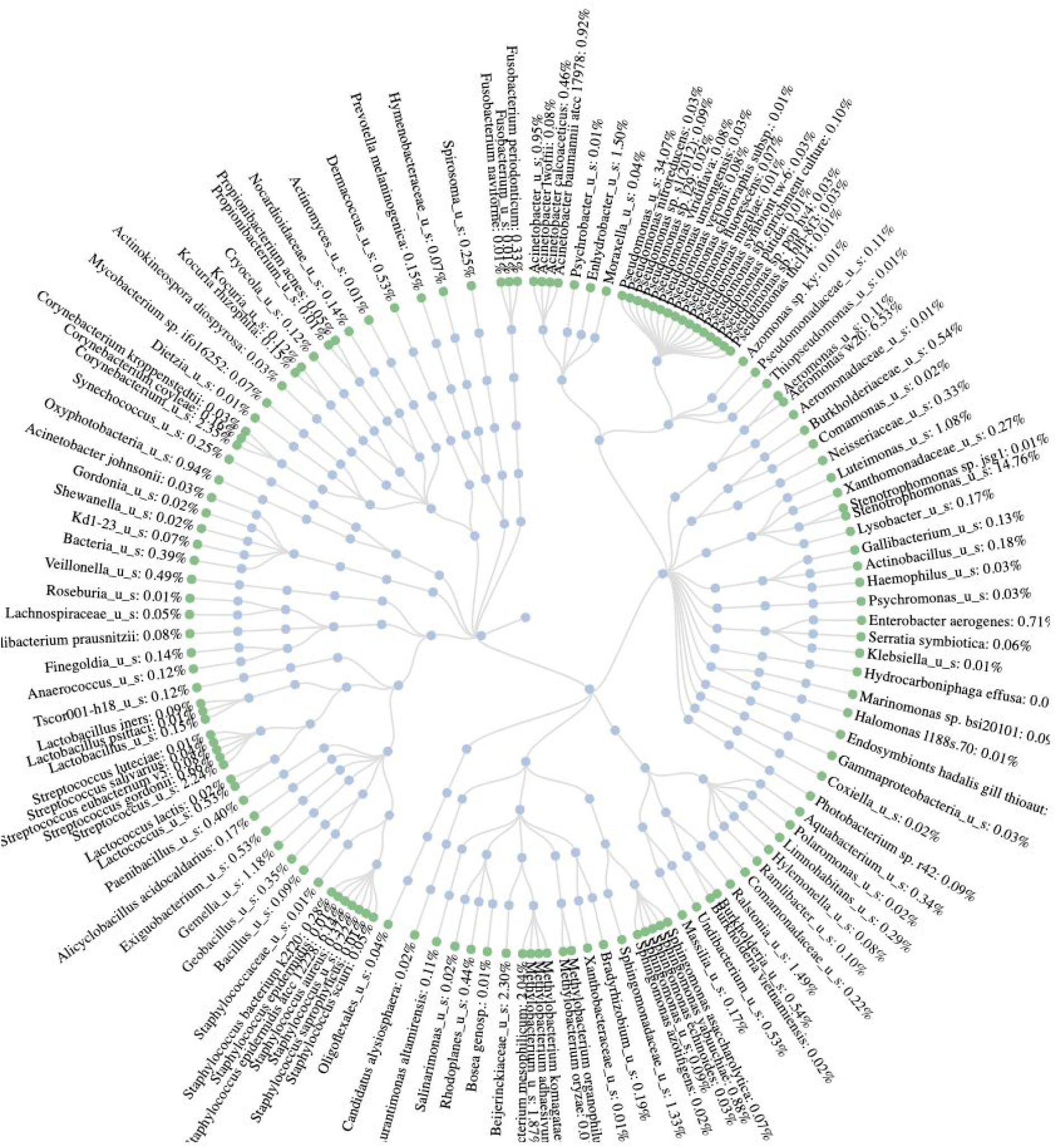
Krona graph of the microbes found in the Forest Hills, Queens, sample using the Soil Kit.

**Figure 3:**
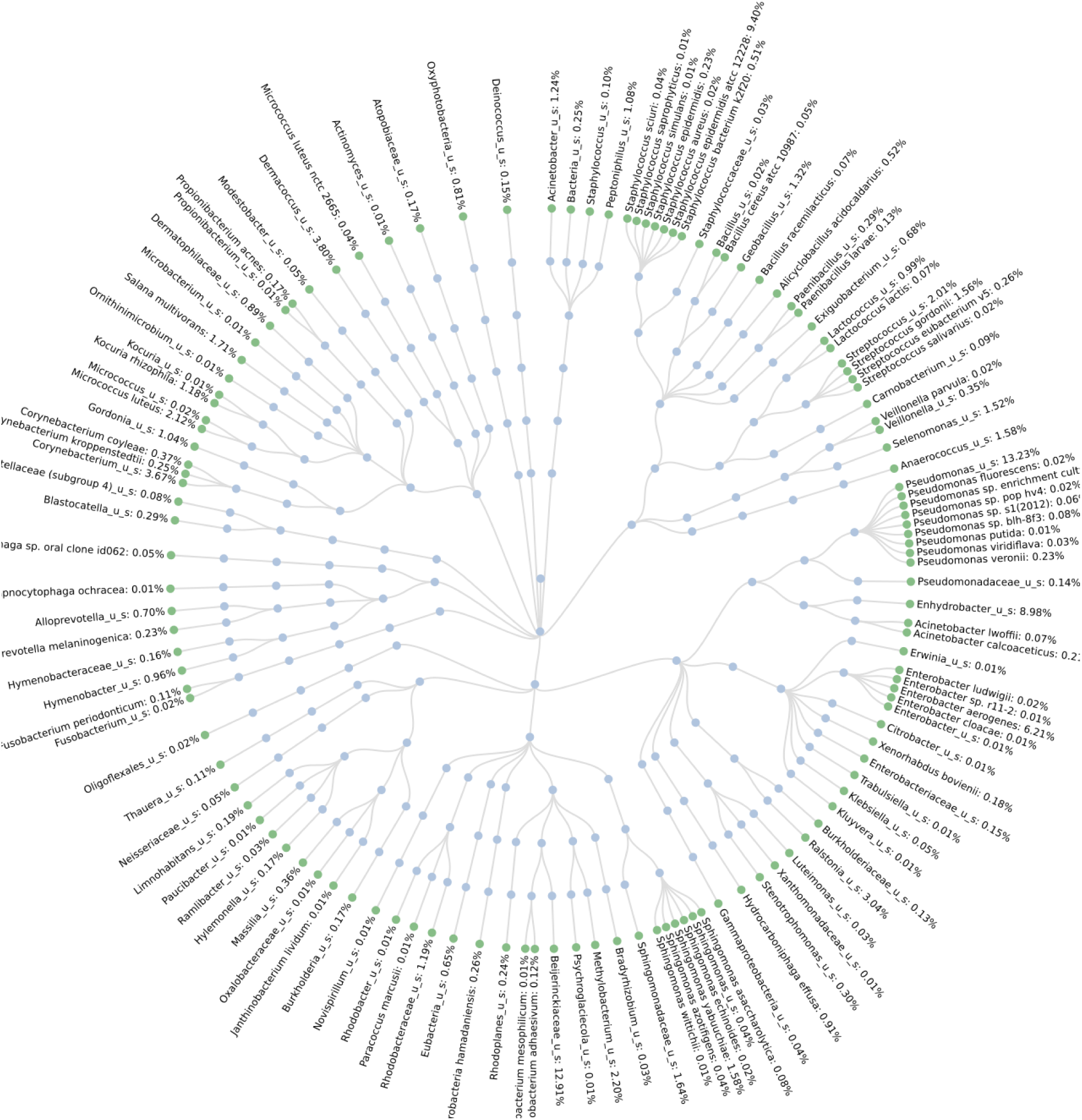
Krona graph of the microbes found in the Rockaway sample using the Soil Kit.

**Figure 4:**
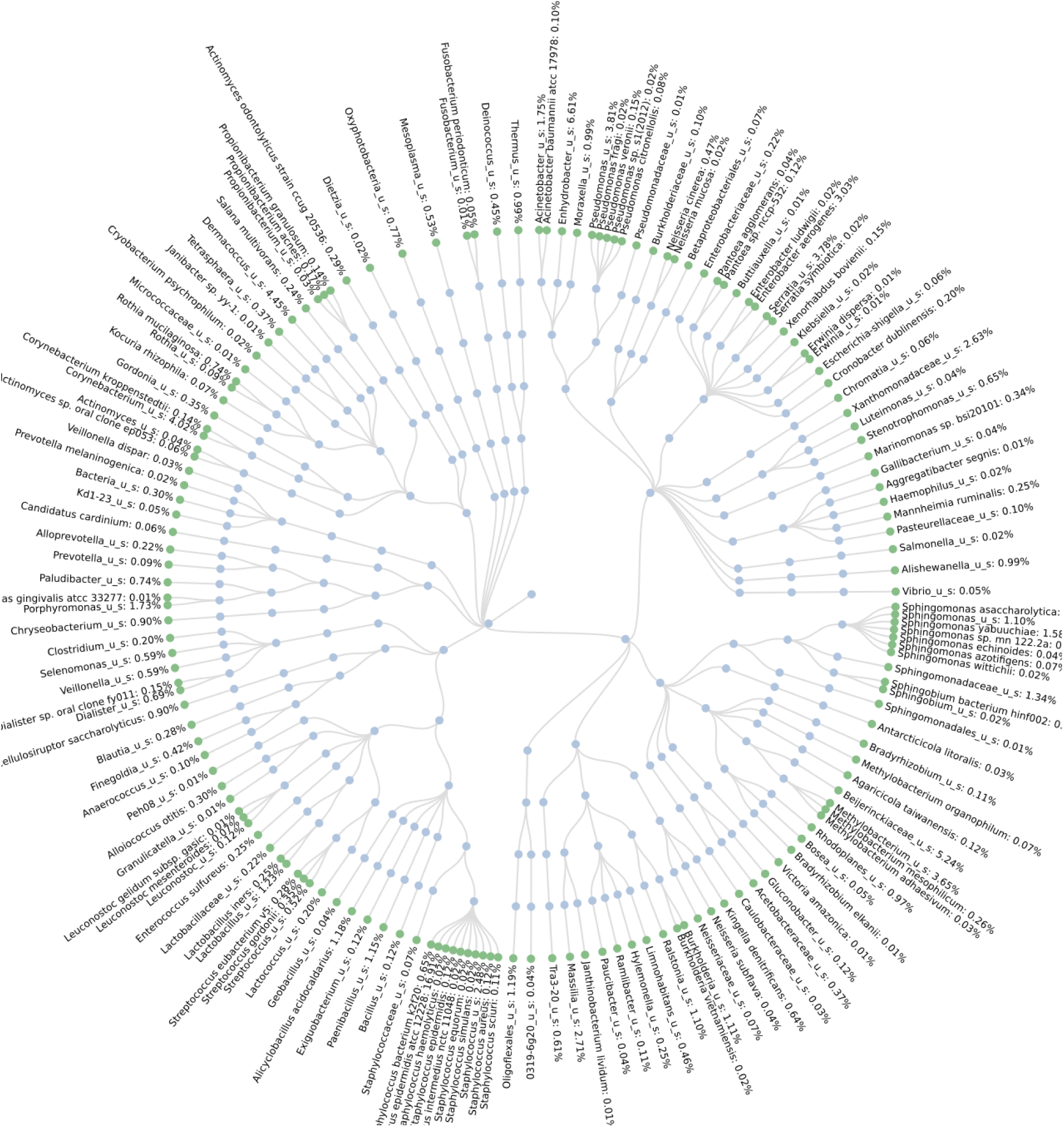
Krona graph of the microbes found in the Westchester County sample using the Soil Kit.

**Figure 5:**
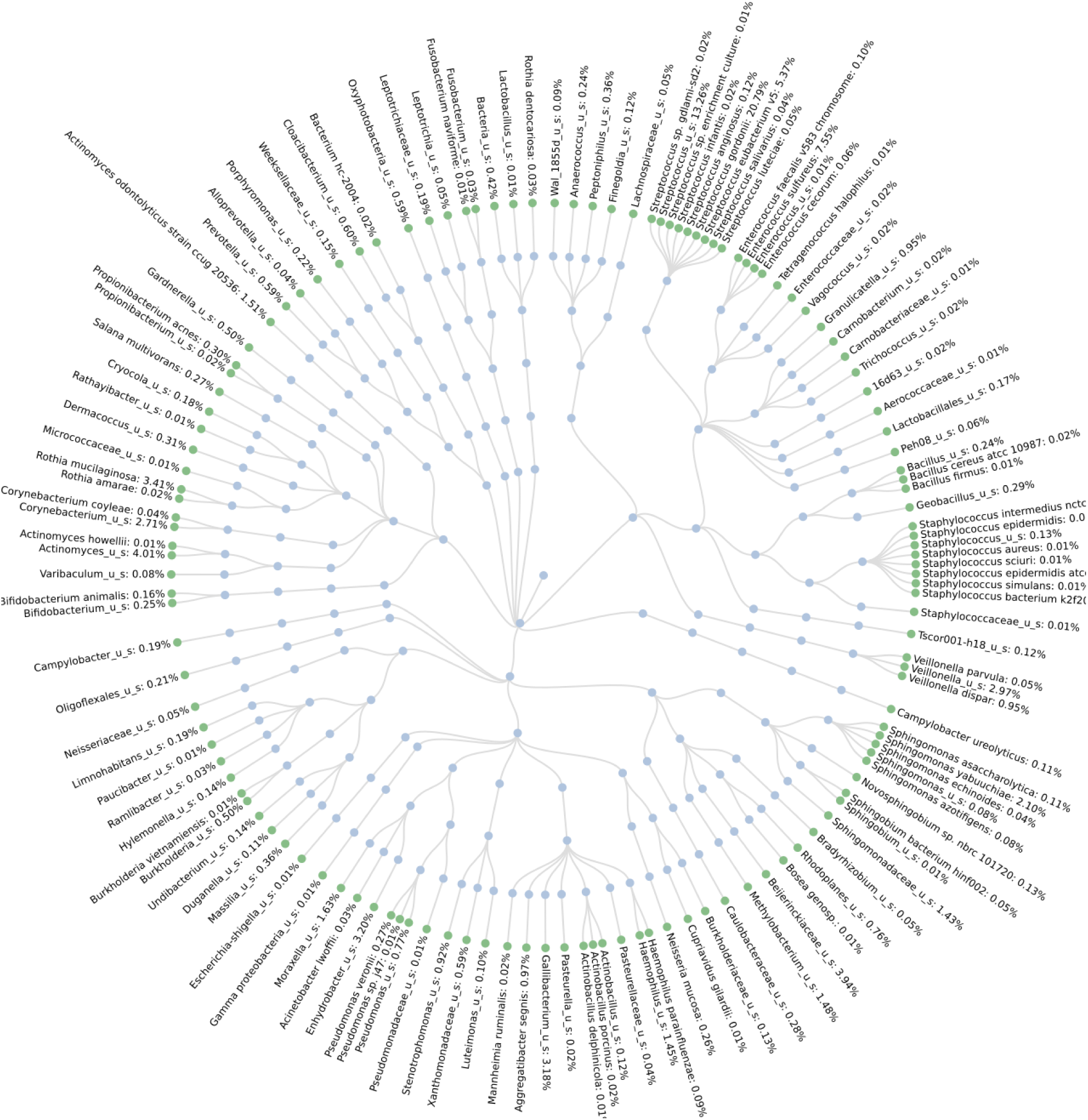
Krona graph of the microbes found in the Madison and 44th Street sample using the Blood and Tissue Kit.

**Figure 6:**
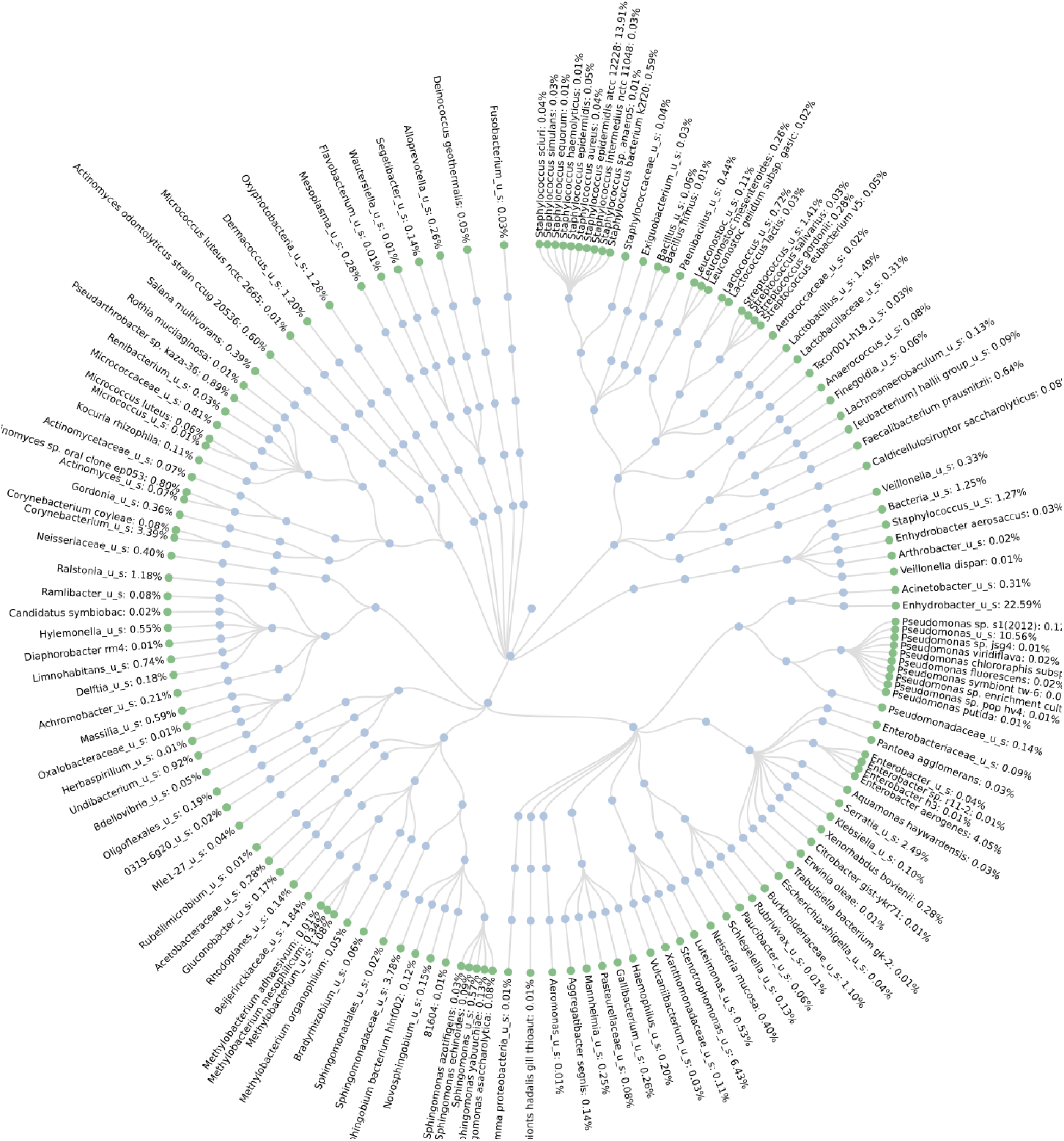
Krona graph of the microbes found in the Harlem sample using the Soil Kit.

**Figure 7:**
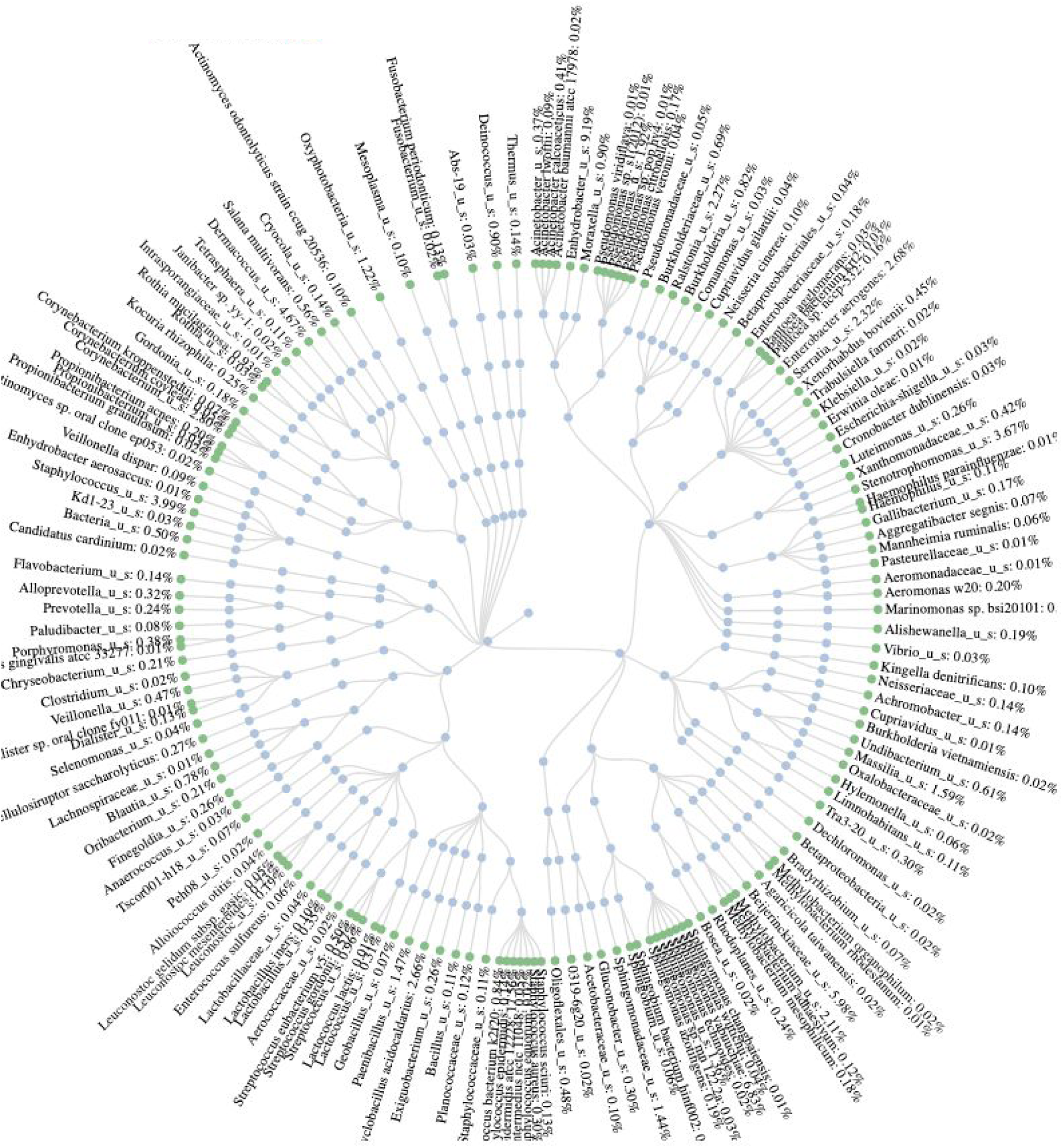
Krona graph of the microbes found in the Store Bought (Pennsylvania) sample using the Soil Kit.

**Figure 8:**
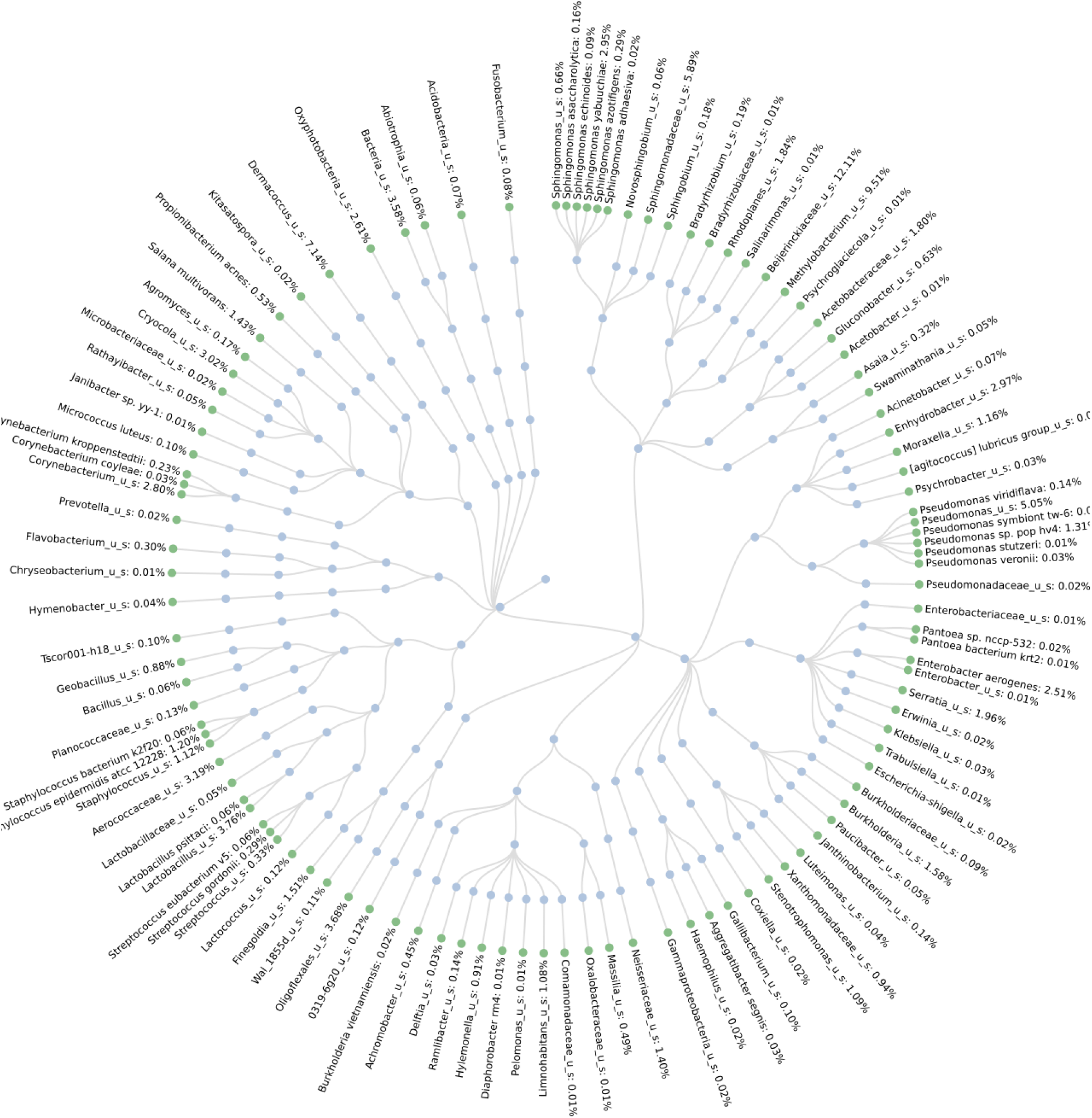
Krona graph of the microbes found in the Store Bought (Pennsylvania) sample using the Blood and Tissue Kit.

**Figure 9:**
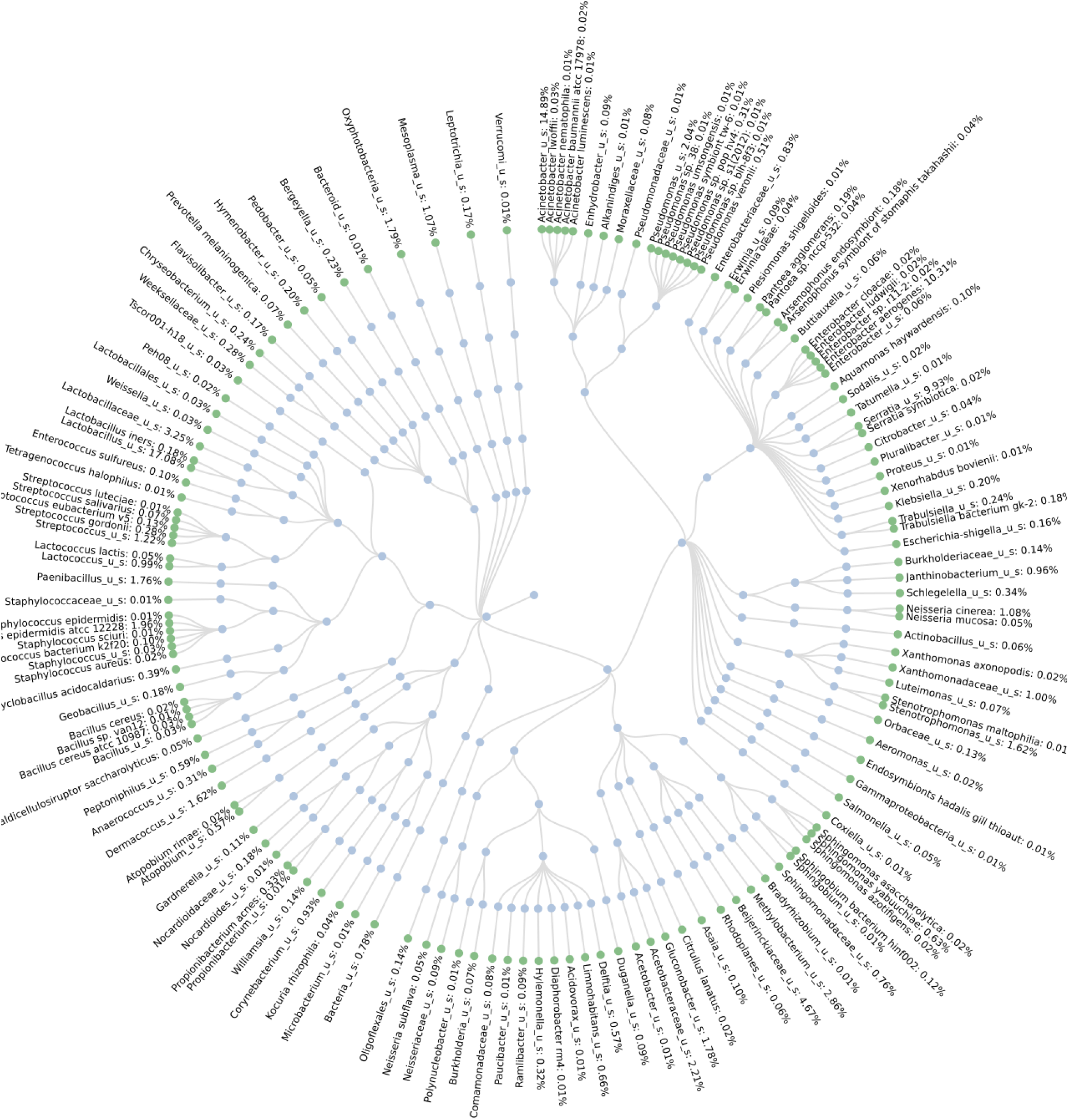
Krona graph of the microbes found in the Harlem sample using the Blood and Tissue Kit.

**Figure 10:**
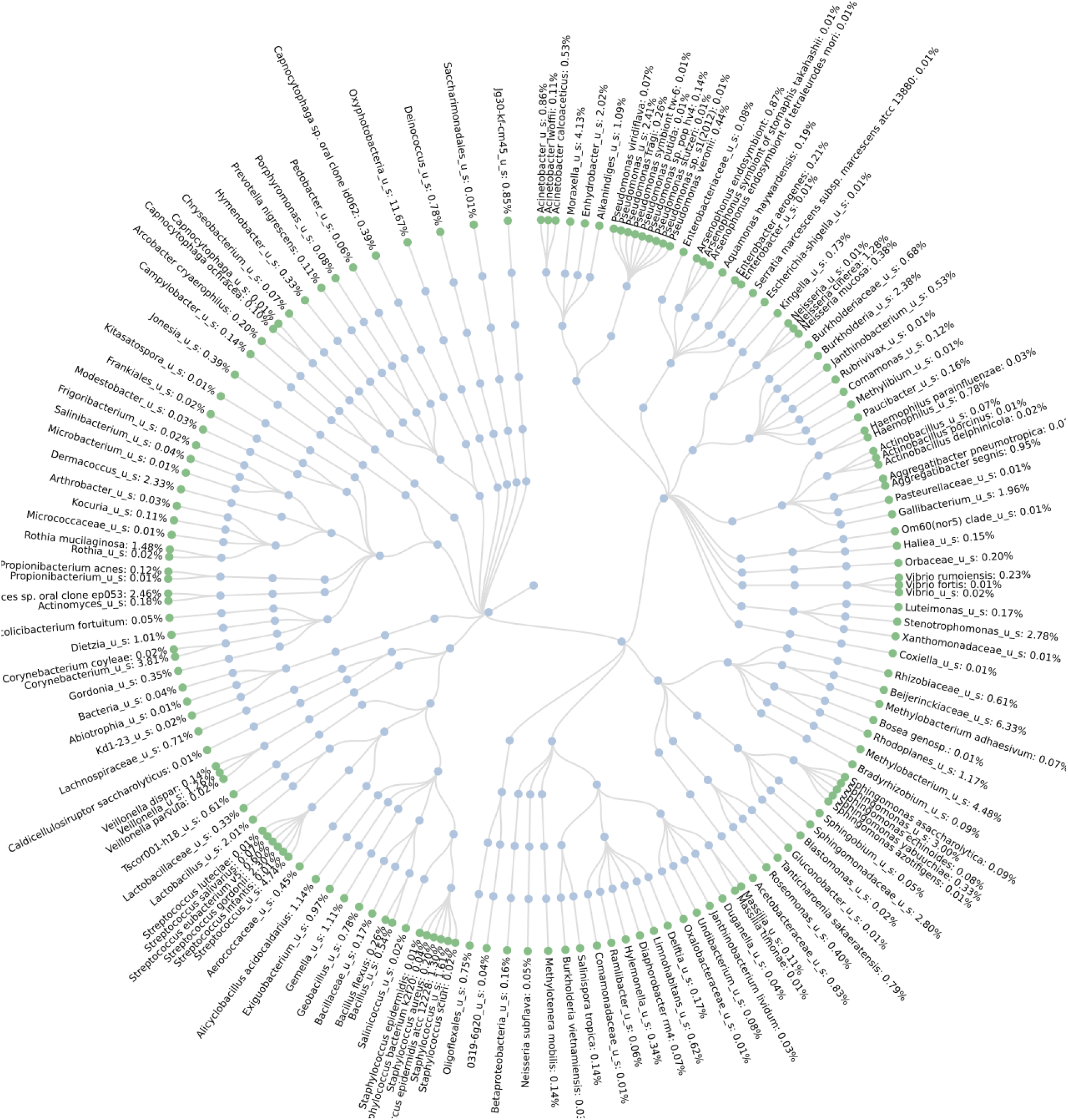
Krona graph of the microbes found in the 86 4th Ave. sample using the Blood and Tissue Kit.

**Figure 11:**
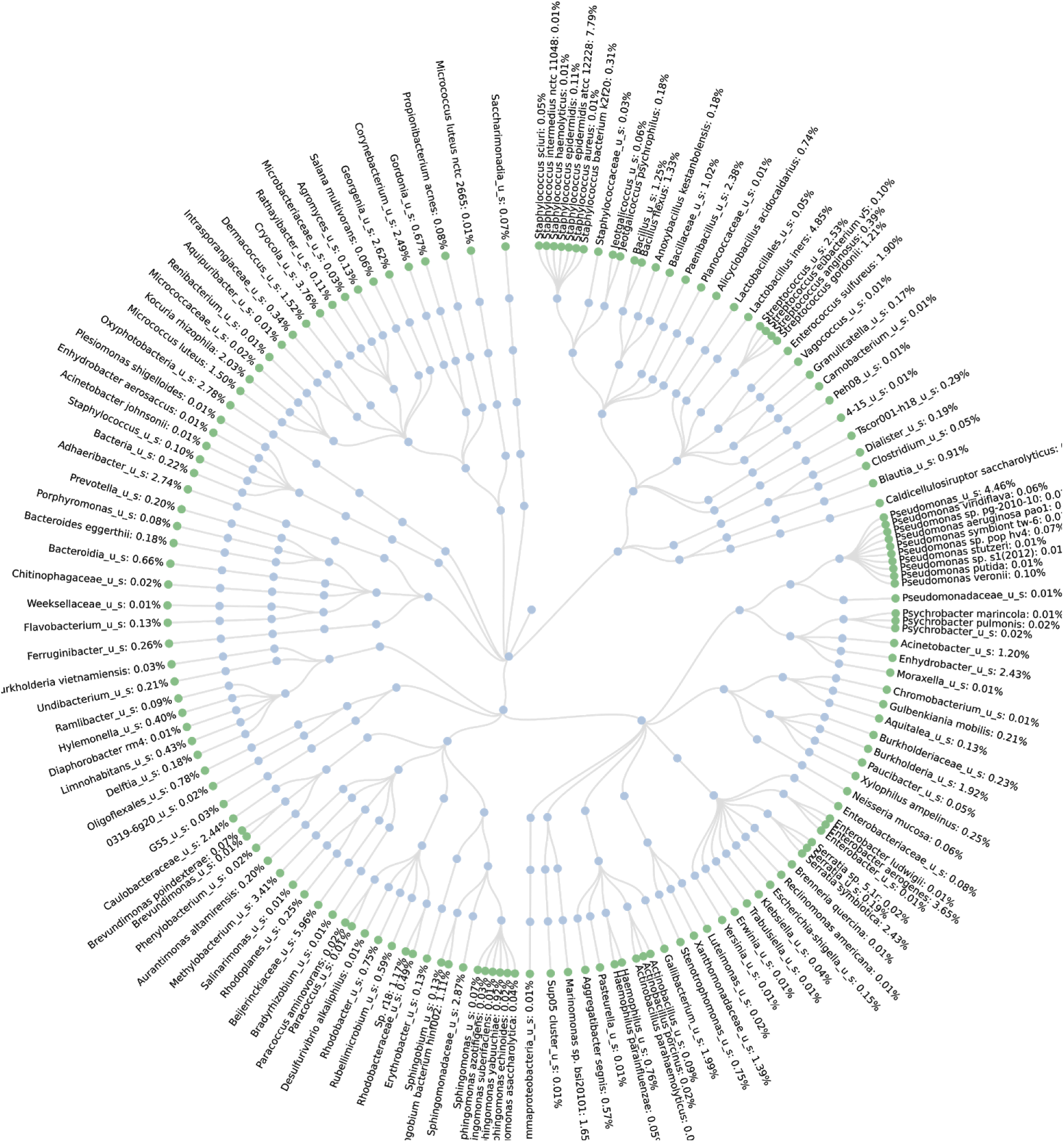
Krona graph of the microbes found in the Westchester County sample using the Blood and Tissue Kit.

**Figure 12:**
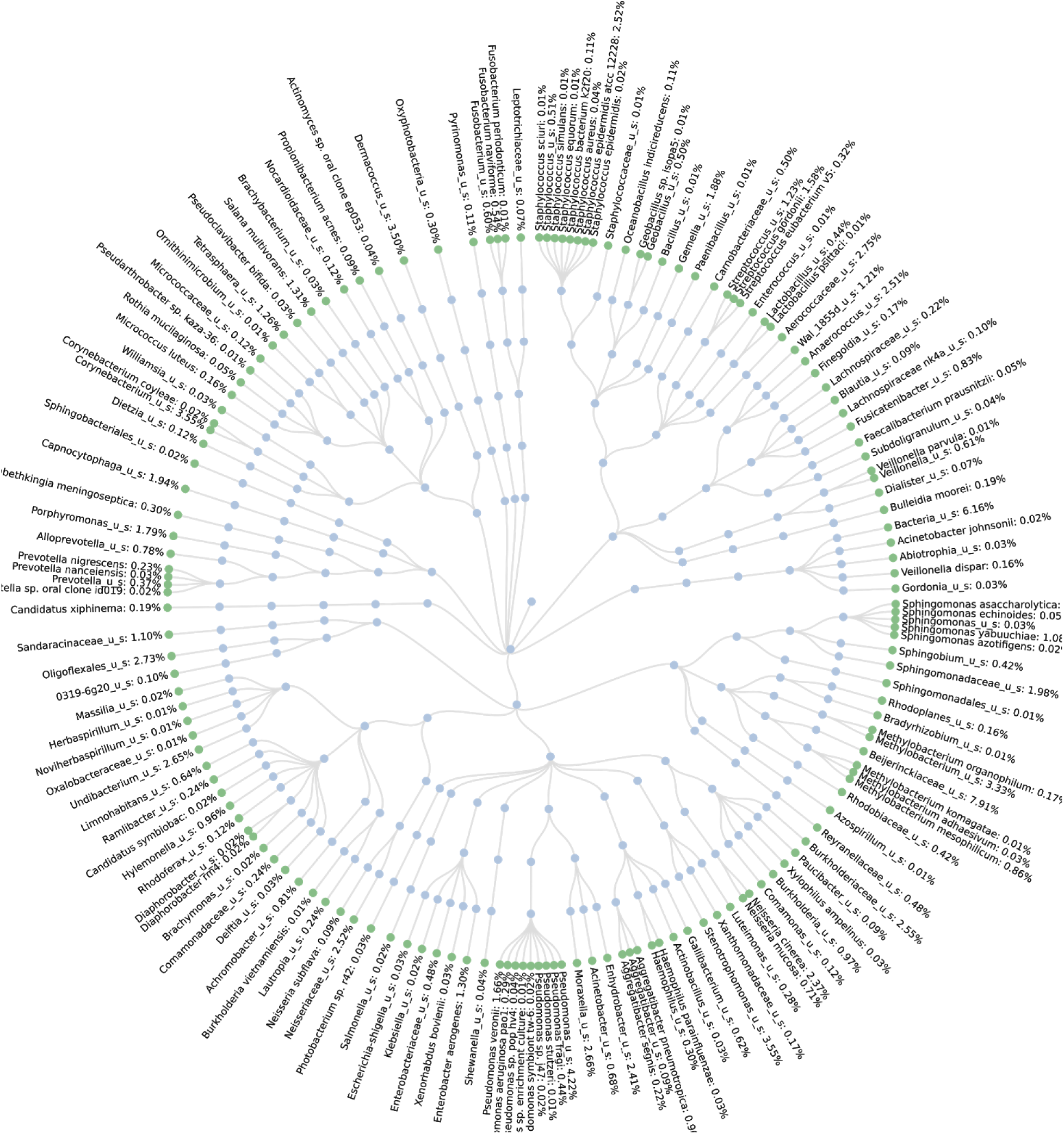
Krona graph of the microbes found in the Financial District sample using Blood and Tissue Kit.

**Table 1:** Percentages of selected microbes identified in samples extracted using the Blood and Tissue Kit.

|  | *Actinobacteria* | *Firmicutes* | *Streptococcaceae gordonii* | *Lactobacillus* | *Cyanobacteria* |
| --- | --- | --- | --- | --- | --- |
| Westchester County | 13.51% | 26.60% | 1.21% | 4.85% | 3.45% |
| 86 4th Ave. | 12.00% | 19.97% | 2.30% | 2.01% | 11.90% |
| Financial District | 10.10% | 17.19% | 1.58% | 0.44% | 0.28% |
| Madison and 44th Street | 14.19% | 55.80% | 10.79% | 0.00% | 0.56% |
| Store Bought / Pennsylvania | 14.72% | 12.00% | 0.29% | 3.76% | 2.50% |
| Harlem | 3.90% | 27.46% | 0.28% | 17.00% | 2.30% |

**Table 2:** Percentages of selected microbes identified in samples extracted using the Soil Kit.

|  | *Actinobacteria* | *Pseudomonas* | *Staphylococcus epidermidis* Strain ATCC 12228 |
| --- | --- | --- | --- |
| Westchester County | 10.79% | 3.81% | 16.91% |
| 86 4th Ave. | 12.40% | 9.72% | 6.92% |
| Forest Hills | 4.17% | 34.07% | 6.14% |
| Rockaway | 15.28% | 13.23% | 9.40% |
| Store Bought / Pennsylvania | 9.40% | 1.92% | 17.56% |
| Harlem | 9.10% | 10.56% | 13.91% |

**Figure 13:**
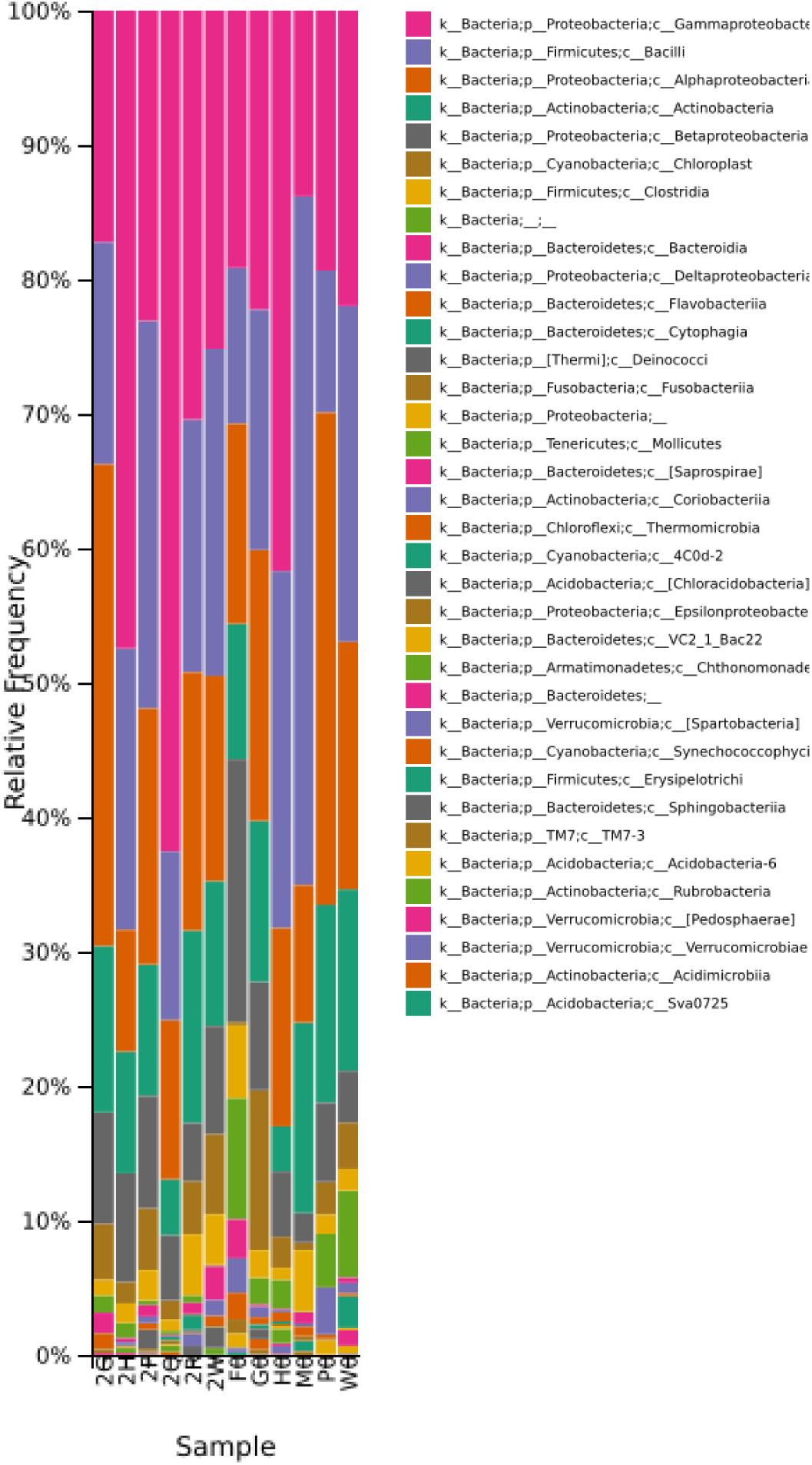
Taxonomic Diversity of the microbes found shown as a bar graph created by the Purple Line on DNA Subway.

The relative abundance of microbes found in the samples. Legend: 2G = 86 4th Ave. - Soil Kit, 2H = Harlem - Soil Kit, 2P = Store Bought / Pennsylvania - Soil Kit, 2Q = Forest Hills, Queens - Soil Kit, 2W = Westchester County - Soil Kit, F0 = Financial District - Blood and Tissue Kit, G0 = 86 4th Ave. - Blood and Tissue Kit, H0 = Harlem - Blood and Tissue Kit, M0 = Madison and 44th - Blood and Tissue Kit, P0 = Store Bought / Pennsylvania - Blood and Tissue Kit, W0 = Westchester County - Blood and Tissue Kit.

278 species were found throughout the different samples and methods, and while the samples contained a variety of similar microbe DNA, there were differences in the abundance rates. Most of the bacteria found are commonly associated with plant growth and have expressed antibiotic properties in humans.

One type of bacteria that was found in all samples was *actinobacteria*, which is a phylum of gram-positive bacteria [8]. As shown in Table 1 and Figure 14, the sequences identified as *actinobacteria* made up 3.94% to 14.72% of sequencing reads using the Blood and Tissue Kit. Using the Soil Kit, Harlem had the lowest percentage, with 3.94% (Fig 6), compared with 14.72% in Pennsylvania (Fig 7). This makes sense because the bees in Pennsylvania most likely came into more contact with soil and agriculture while foraging than those in New York City. *Actinobacteria* commonly breaks down dead organic matter [8], is found in the guts of bees, and contributes to honey’s medicinal properties [9].

**Figure 14:**
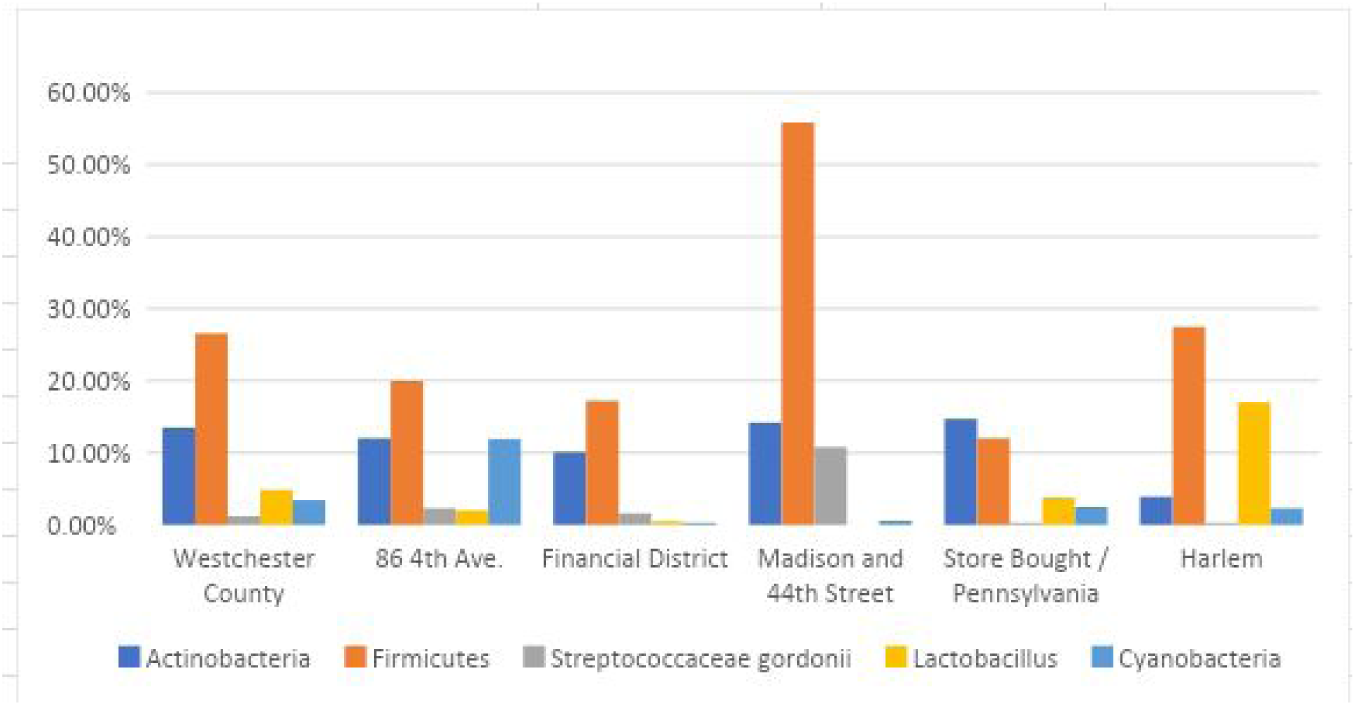
Percentages of selected microbes identified in samples extracted using the Blood and Tissue Kit.

Another aspect of honey that the microbes influence is its taste. *Methylobacterium* was found in all of the samples in a variety of quantities, and studies suggest that *Methylobacterium* is responsible for the flavor of strawberries [10]. It is possible that honey with a larger percentage of *Methylobacterium* would taste more like strawberries.

### Results using the Blood and Tissue Kit

*Firmicutes* were also found. They are gram-positive and comprise the largest portion of the human and mouse gut microbes [11]. Samples that were collected closer to the center of Manhattan had a higher percentage of *firmicutes*. Using the Blood and Tissue Kit, *firmicutes* made up only 17.19% of the sample taken from the Financial District (Fig 12). The highest concentration of *firmicutes* was found on Madison and 44th Street, at 55.82% (Fig 5), possibly due to its higher concentration of people and rodents.

Figure 5 shows that 20.79% of the sample taken from Madison and 44th Street was *Streptococcaceae gordonii*, while the rest of the samples contained between 0.28% and 2.30% (Table 1, Fig 14). *S. gordonii* inhabit the human mouth, skin, intestines, and upper respiratory tract [12]. Even though they are known to form dental plaque on teeth [12], honey has been shown to not cause dental erosion [13]. This microbe relies on sugar for energy [12], suggesting that the honey from Madison and 44th Street contained a higher concentration of sugar compared with the other samples. Another possibility is that the bees had more access to *S. gordonii* at this location than the others.

From the Harlem sample, 17.08% of the sequence was from the genus *Lactobacillus* (Fig 9). For reference the other samples contained between 0% and 2.01% (Table 1, Fig 14). Much research has been done into the probiotics properties of *Lactobacillus* as well as its ability to treat and prevent diseases [14].

One surprising microbe that was found was the photosynthetic bacteria *cyanobacteria.* It represented 11.38% (Fig 13) of the sequences in the sample from 86 4th Ave compared with only 1.35% to 3.5% (Fig 13) of the other samples. C*yanobacteria* has anti-inflammatory and antimicrobial properties, however under certain conditions *cyanobacteria* can produce cyanotoxins that are harmful to humans [15]. Cyanobacteria can also damage honey bee populations by increasing the mortality rate and reducing their ability to learn odors [16]. Through trophallaxis, these toxins are transmissible from bee to bee, and can infect the entire hive [16]. This is another theorized cause of Colony Collapse Disorder. Studies suggest that people living near lakes with high levels of *cyanobacteria* are more likely to develop Amyotrophic Lateral Sclerosis (ALS) [17]. The bees in this hive appear to be foraging someplace with a higher amount of *cyanobacteria* than the other locations thus giving an idea of what the bees and possibly people are exposed to.

### Results using the soil sample kit

*Pseudomonas* was found to make up 34.07% of the sample from Forest Hills, Queens (Fig 2), while the rest of the samples only had between 1.92% and 13.23% (Table 2, Fig 15). *Pseudomonas* is a gram-negative *gammaproteobacteria* and is commonly found in soil, vegetation and water [18]. This implies that the area surrounding the hive is more densely populated by vegetation than the other locations.

**Figure 15:**
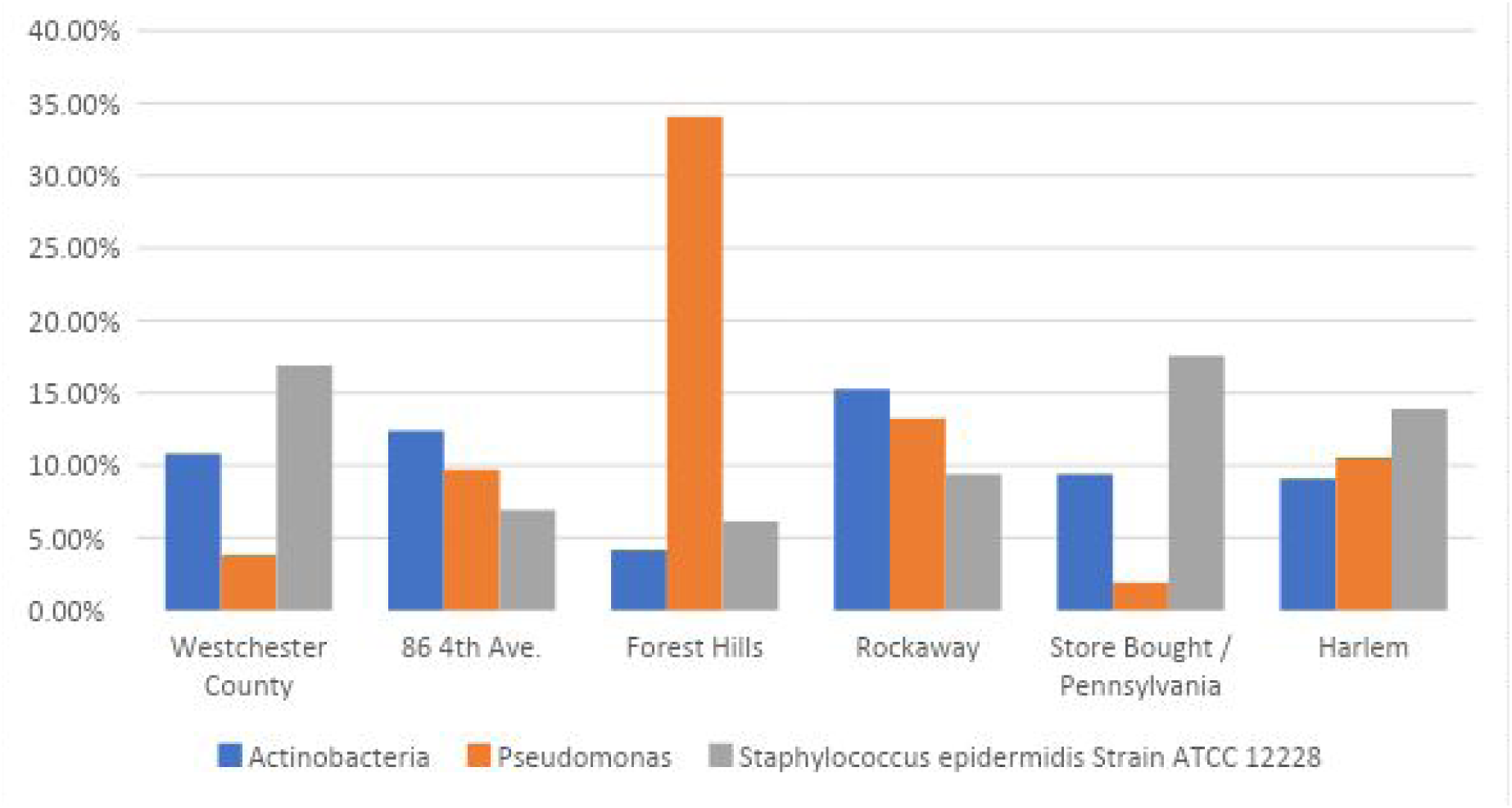
Percentages of selected microbes identified in samples extracted using the Soil Kit.

*Staphylococcus epidermidis* is a gram-positive facultative anaerobic bacteria found on human skin [19]. The samples from Pennsylvania and Westchester County had the highest percentage of *S. epidermidis*, at 16.91% (Fig 7) and 17.58% (Fig 5), respectively, compared with the other samples, which had between 6.14% and 13.9% (Table 2, Fig 15). This is surprising because *S. epidermidis* is typically found on skin, and the highest concentration is in areas with a lower concentration of people, as opposed to the densely populated Manhattan. We cannot rule out contamination by human contact in the collection or processing of these samples.

## Discussion

The honey bee gut microbiome has been shown to contain *Lactobacilli*, which was also found in late larval stages, suggesting that the Harlem sample had a greater abundance of older larvae and adults compared with the other samples [20]. *Firmicutes, gammaproteobacteria*, and *actinobacteria* were also found in the current study and another study that looked at the relationship between the microbes found, the geographical location, and the age of the bees [20]. One thing we found that the other study did not was the presence of *cyanobacteria.*

In a study published by the Journal of Global Infectious Diseases, *Pseudomonas* was also found in the honey samples [21]. That experiment examined the antimicrobial properties of honey and its ability to kill *P. aeruginosa* [21]. These cited experiments used honey collected from different parts of the world, while we used honey from New York City, Westchester County and Pennsylvania. Despite the geological differences, the results of each study demonstrate the consistent presence of the microbes *Lactobacillus, firmicutes, gammaproteobacteria, actinobacteria*, and *Pseudomonas* species.

This research provides a way to look at individual neighborhoods in New York City and compare them with each other while also providing an understanding of the microbes of the areas. The microbes we found are from the bee’s gut microbiome, the area surrounding the hive, and areas where the bees are pollinating and retrieving the honey. By tracking this we can determine what the bees, and hence possibly the humans, in the respective neighborhoods are coming in contact with. Colony Collapse Disorder is causing the bee populations to decline at a rapid rate, and by studying what the bees interact with in their areas, one could find microbes that harm the bees and prevent their further destruction; bees could also act as collectors of microbes relevant to human or overall environmental health, acting as “monitors” of microbial diversity in their ranges.

## Conclusion

The honey samples were all distinct and contained a wide variety of microbes clearly influenced by their locations. All taxonomic plots created from the data are unique; however, the differences found in the non-city samples were not greater than those found comparing the city samples with each other. This study was limited in scope due to the short time frame of the student-driven research project. Our ability to sequence replicates was also limited, and future research should focus on repeating and confirming these findings, as well as performing statistical analyses on the differences between samples. The research is consistent with other studies and reinforces the similarities of honey across geographic locations. The microbes found also contribute to honey’s antibiotic properties. This study shows that honey produced by urban beekeeping is not drastically different than honey from rural beekeeping. This is despite the fact that there are many differences between the environments such as the people and vegetation. Results from all locations were different, yet many of the microbes found were the same between the samples done in this experiment and others that used honey from around the world.

## Acknowledgments

Thank you to DNALC NYC Partner Member Program for all of its support and funding. *DNA Subway* was developed under CyVerse (formerly iPlant Collaborative), a cyberinfrastructure project supported by the National Science Foundation (DBI-0735191, DBI-1265383, and DBI-1743442). Development of the Purple Line of *DNA Subway* was supported in part by NIH SEPA Microbiome BD2K supplement 3R25OD016511-03S1 and grants from the Richard Lounsbery Foundation. We would like to thank Grace Church High School for all its support, in addition to Dr. Sibylle Brenner and Cynthia Jackson for all of their help. Thank you to Andrew Cote for contributing the honey samples. We thank Dr. Larry Weiss for providing the inspiration for this project and guidance throughout. We thank Nur Hasan and Brian Fanelli for creating the krona graphs through Cosmos ID. Lastly, thank you to Aliya Weiss and Fiona Humphrey for editing this paper.

